# Gut microbiota analysis of the western honeybee (*Apis mellifera* L.) infested with the mite *Varroa destructor* reveals altered bacterial and archaeal community

**DOI:** 10.1101/2022.04.20.488909

**Authors:** Minji Kim, Woo Jae Kim, Soo-Je Park

## Abstract

The western honeybee, *Apis mellifera* L., is a crop pollinator that makes royal jelly and other hive products. However, widespread concerns arise about opportunistic diseases (e.g., bacteria, fungi, or mites) or chemicals that have an effect on the health and number of colonies, as well as their activity. The relationships between the gut microbiota and its host are currently being researched extensively. The effects of *Varroa destructor* infection on the gut microbial community, in particular, have received little investigation. This work utilized amplicon sequencing of the bacterial and archaeal 16S rRNA genes to assess the bacterial and archaeal communities of adult bee groups (healthy and affected by *Varroa* designed in NG and VG, respectively) and larvae from *Varroa destructor*-infected hives. Our results suggest that the genus *Bombella* was substantially dominant in larvae, while the genera *Gillamella*, unidentified *Lactobacillaceae*, and *Snodgrassella* were significantly dominant in adult bees. NG and VG, on the other hand, did not differ statistically significantly. The PICRUSt study revealed a significant difference in the KEGG classifications of larvae and adult bee groups. A greater number of genes involved in cofactor and vitamin production were identified in larvae. Additionally, despite the complexity of the honeybee’s bacterial community, all groups exhibited a straightforward archaeal community structure. Surprisingly, methanogen was detected in low abundance in the microbiota of honeybees. In summary, larvae and adult bees infected with *Varroa destructor* exhibit altered gut microbiota composition and function.

## Introduction

The next-generation sequencing method has considerably expanded the exploration of the microbiome and its contribution to the host to vertebrate (e.g., human) or plant as a leading scientific field. In particular, the studies for microorganism and/or microbial communities colonized in the host gastrointestinal tract have improved our understanding and knowledge of the ecological and functional roles in its gut environments. The gut microbial communities (i.e., microbiota) are recognized as being critical for the survival of a wide variety of host organisms.. Nonetheless, the intricate co-evolutionary relationships between microorganisms and hosts, as well as the potential/fundamental properties of gut microorganisms, remain largely unknown.

The western honeybee, *Apis mellifera* L, is a key player as a pollinator species for natural ecosystem and agricultural production (1). *Ap. mellifera* is critical because it contributes to the pollination of over 90 percent of key commercial crops in the United States of America (2). Among the crops, blueberries and cherries are 90% depending on their pollination activity. The entire of the almond crop depends on the *Ap. mellifera* for pollination at bloom time (American Beekeeping Federation, http://www.abfnet.org). Additionally, honey and beeswax production are valuable commodities to humans and contribute to the economic value of honey bees. Despite their critical contributions to human consumption, the colony sustainability of honeybees has been changed by modern agricultural practices (e.g., use of pesticides and agrochemicals), pathogen exposure, or environmental changes (e.g., global warming) (3-7).

Recent research indicates that *Ap. mellifera* gut dysbiosis induced by antibiotic and microplastics exposure may impair their activity (8-11). Additionally, these researches indicated a possible hazard to public health posed by opportunistic human infections that acquired antibiotic resistance genes from drug-treated honeybees. Numerous prudential applications, including probiotics, have been made to honeybee colonies in response to the situation (12-14). *Lactobacilli* spp., in particular, enhances honeybee activity, stress control, and queen brood production (13, 15-18). Apart from the factors discussed above, the resulting gut microbiota may be influenced by infectious disease transmitted by viruses, microsporidian, or mites, which may have an effect on honeybee health, activity, and population (19-22). To summarize, examining the *Ap. mellifera* gut microbiota as a novel experimental paradigm is crucial for future research on the human gut microbiota (23-26).

Numerous studies have examined the compositions or changes in the honeybee gut microbiota of different bee classes (i.e., forage, nurse, or queen) (27-30). Until now, the majority of studies on the microbial community have concentrated on the entire body of honeybees or mites (31-35). To our knowledge, the gut microbiota of larvae and adult bees infected with *Varroa destructor* has not been comprehensively investigated.

The purpose of this study was to i) characterize and compare the archaeal and bacterial community structures found in larvae and adult honeybees from the *Varroa*-infested hives; and ii) estimate differences in putative functional roles based on the microbial compositions of larvae and adult bees. Our findings may contribute to a better understanding of the interaction between microbiota and honeybees, as well as their functional roles in the honeybee gut environment.

## Materials and methods

### Sample collection

We surveyed beehives afflicted with the ectoparasitic mite *Varroa destructor* from beekeeping farms on Jeju Island toward the end of May 2021, practically to the end of the full blooming period. Finally, two larvae and fourteen adult bees were collected from two western honeybee (*Ap. mellifera* L.) apiaries, one without varroa (NG, n=9) and one with varroa (VG, n=5). Two *Varroa*-infected apiaries were discovered by an experienced beekeeper, who confirmed the deceased *Varroa* and clinical symptoms (e.g., shortening of the wing) with his naked eyes. After harvesting samples into the sterile falcon tube, they were immediately kept at a low temperature (4°C) using an icepack and sent to the laboratory for further processing.

### DNA extraction and amplicon sequencing

Total genomic DNA (gDNA) was extracted from isolated adult bee guts or whole-body for larva using a QIAamp PowerFecal Pro DNA Kit (Qiagen). Gut tract from each surface-sterilized adult bee was dissected in sterilized phosphate buffered solution (PBS, pH 7.4) under anatomical microscope. The quality and quantity for the extracted gDNA were estimated by a DS-11 Plus Spectrophotometer (DeNovix, Inc., Wilmington, DE) and confirmed agarose gel (1.5% w/v) electrophoresis. The gDNA samples were frozen at -20°C for further experiment.

To obtain the amplicon for bacterial and archaeal 16S rRNA gene, we conducted PCR on an Illumina platform according to our previously describe studies (36). Briefly, total 20 µl of PCR mixture was prepared as follows: 10 µl of Solg™ 2x EF-Taq PCR Smart mix (Solgent, South Korea), 1 µM primer set (final conc.), and about ∼5 ng of template gDNA. The procedures for thermal amplification were as follows: an initial denaturation step at 95°C for 5 min; followed by 30 cycles of 95°C for 30s, 55°C for 30s and 72°C for 40s, ended with a final extension step at 72°C for 7min. The sequences of the primer sets were targeted to the V4-V5 hyper-variable region of 16S rRNA gene for Bacteria (515F, 5’-GTGCCAGCMGCCGCGGTAA-3’ and 907R, 5’-CCGTCAATTCCTTTGAGTTT-3’) and Archaea (519F, 5’-CAGCCGCCGCGGTAA-3’ and 915R, 5’-GTGCTCCCCCGCCAATTCCT-3’). PCR amplified products were visualized by 1.5% (w/v) agarose gel electrophoresis for amplified size confirmation. Then, the amplicons were purified with the Monarch® PCR & DNA Cleanup Kit (NEB). High-throughput sequencing was performed by Novogene using the Illumina NovaSeq PE250 system (Illumina, Inc.), according to the manufacturer’s instructions.

### Data analysis and statistics

Sequencing data was analyzed using the standard operating procedure (SOP) described on Mothur (version 1.46.1) website (https://mothur.org/wiki/miseq_sop/). All raw reads were obtained after trimming the barcode and primer sequences. The trimmed paired-end reads were merged supplied in Mothur program. Subsequently, high-qualified merged reads were obtained by filtering and chimeric sequences were removed using *chimera*.*vsearch* command. Also, to increase the analysis quality, the qualified-read sequences were unknown and non-microbial sequences (e.g., chloroplast, mitochondria, and eukaryote), were discharged. Then, the sequences were assigned to operational taxonomic units (OTUs) at 97% sequence similarity and a representative sequence was selected from each OTU. The bacterial and archaeal sequences were aligned and classified to a reference database (version silva.nr_v138.1) provided Mothur website (https://mothur.org/wiki/silva_reference_files/). Alpha-diversity indices (i.e., Chao1 nonparametric richness, Shannon, inverse-Simpson, and Good’s coverage) and beta-diversity [i.e., unweighted pair group method with arithmetic mean (UPGMA) clustering and principle coordination analysis (PCoA)] were estimated by the Mothur package. Analysis of molecular variance (AMOVA) was performed to compare the diversity indices for microbial community between two groups. Unless otherwise state, the proportion of total sequences representing each sample or group (combined from same experimental sample) was calculated. Heatmap was generated using R package (gplots). The differences in taxa between the two groups was carried out the linear discriminant analysis effect size (LEfSe) in an on-line interface using default parameters (threshold of > 2.0 for the logarithmic LDA score).

Phylogenetic investigation of communities by reconstruction of unobserved states (PICRISt2) was conducted to predict putative functional profiles based on the microbial community. Bacterial functional profiles were annotated using the Kyoto Encyclopedia of Genes and Genomes (KEGG) pathways.

## Results

### General features of bacterial diversity of honeybee gut microbiota

A total of 2,531,114 raw reads were acquired from two larvae (designated as L) and fourteen adult bees [attached or unattached *Varroa* designated as varroa group (VG) or non-varroa group (NG), respectively] and each sample was sub-sampled by 20,000 reads. The number of reads used for sub-analysis processing varied between 2041 and 3421 reads per sample.

The diversity indices were estimated using the qualified and subsampled reads. The detailed diversity values for larva and adult bee were shown in Table 1. Larvae had a significantly different gut microbiota than NG (ANOVA, p=0.02) and VG (p=0.009) (sFig. S1). Surprisingly, there was no significant difference in microbiota between NG and VG (ANOVA, p=0.292) (sFig. S1). The results for the number of the estimated OTUs (Chao1) indicates that NG and VG are higher than L group (Kruskal-Wallis, p=0.059 and 0.053, respectively). However, statistical analysis of the diversity indices reveals no substantial inter-group differences. Furthermore, the Good’s coverage (avg. 65.8 percent) indicated that microbial diversity did not achieve a horizontal asymptote, implying that the sequencing effort did not saturate diversity (Table 1).

**Table 1.**
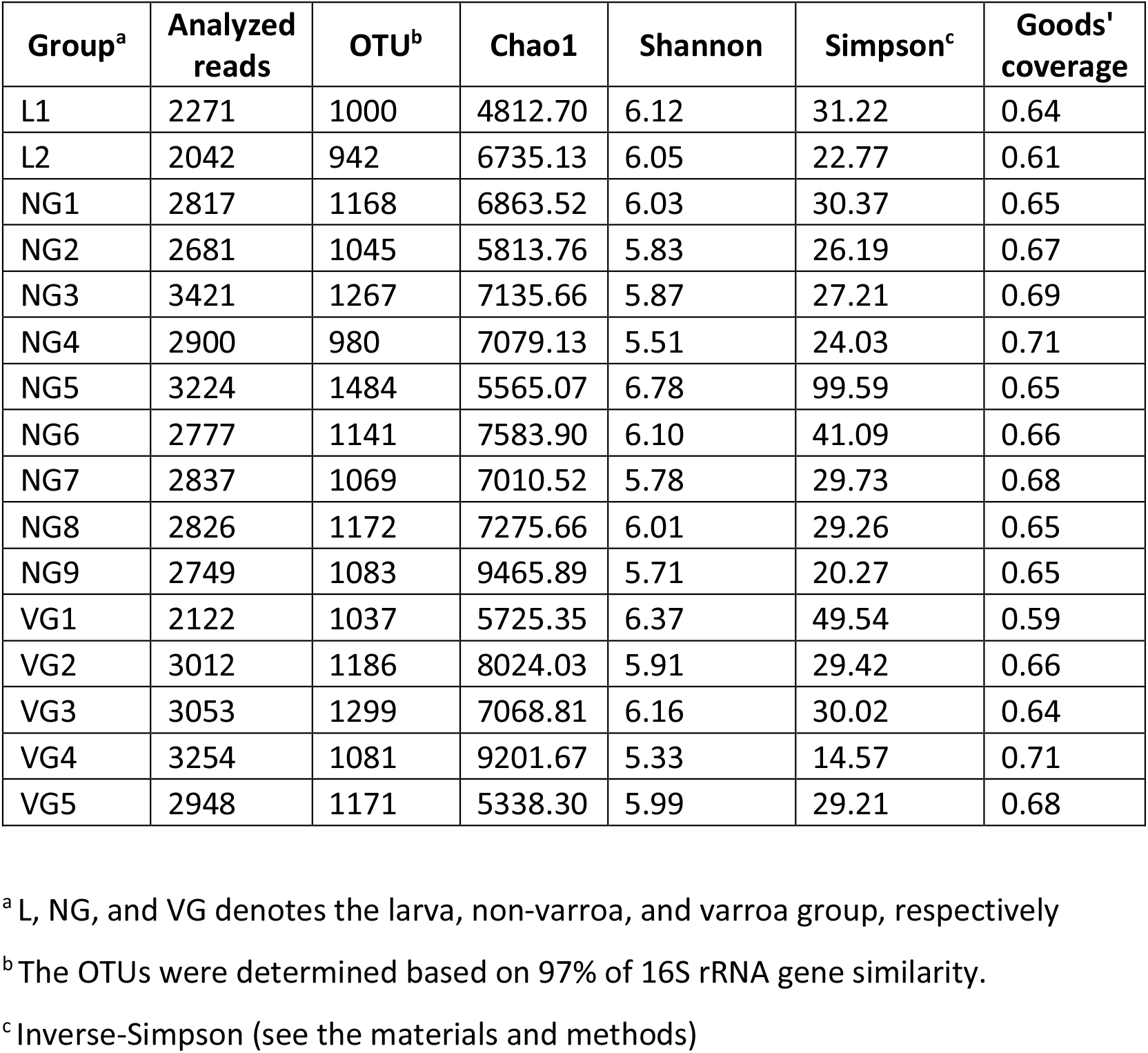
An overview of estimates of read sequence diversity and phylotype coverage of the NovaSeq data generated from the larva and adult bee samples. The diversity indices and richness estimators were calculated using mothur software. Diversity was estimated using operational taxonomic units (OTUs) and was defined as groups with ≥ 97% sequence similarity.

| Group<sup>a</sup> | Analyzed reads | OTU<sup>b</sup> | Chao1 | Shannon | Simpson<sup>c</sup> | Goods' coverage |
| --- | --- | --- | --- | --- | --- | --- |
| L1 | 2271 | 1000 | 4812.70 | 6.12 | 31.22 | 0.64 |
| L2 | 2042 | 942 | 6735.13 | 6.05 | 22.77 | 0.61 |
| NG1 | 2817 | 1168 | 6863.52 | 6.03 | 30.37 | 0.65 |
| NG2 | 2681 | 1045 | 5813.76 | 5.83 | 26.19 | 0.67 |
| NG3 | 3421 | 1267 | 7135.66 | 5.87 | 27.21 | 0.69 |
| NG4 | 2900 | 980 | 7079.13 | 5.51 | 24.03 | 0.71 |
| NG5 | 3224 | 1484 | 5565.07 | 6.78 | 99.59 | 0.65 |
| NG6 | 2777 | 1141 | 7583.90 | 6.10 | 41.09 | 0.66 |
| NG7 | 2837 | 1069 | 7010.52 | 5.78 | 29.73 | 0.68 |
| NG8 | 2826 | 1172 | 7275.66 | 6.01 | 29.26 | 0.65 |
| NG9 | 2749 | 1083 | 9465.89 | 5.71 | 20.27 | 0.65 |
| VG1 | 2122 | 1037 | 5725.35 | 6.37 | 49.54 | 0.59 |
| VG2 | 3012 | 1186 | 8024.03 | 5.91 | 29.42 | 0.66 |
| VG3 | 3053 | 1299 | 7068.81 | 6.16 | 30.02 | 0.64 |
| VG4 | 3254 | 1081 | 9201.67 | 5.33 | 14.57 | 0.71 |
| VG5 | 2948 | 1171 | 5338.30 | 5.99 | 29.21 | 0.68 |

<sup>a</sup> L, NG, and VG denotes the larva, non-varroa, and varroa group, respectively
<sup>b</sup> The OTUs were determined based on 97% of 16S rRNA gene similarity.
<sup>c</sup> Inverse-Simpson (see the materials and methods)

Although the diversity indices for gut microbiota did not show statistically significant differences between intergroups, the results of the unweighted pair group method with arithmetic mean clustering (UPGMA) and principal coordinates analysis (PCoA) indicated that the honeybee samples were clearly divided into two groups at the OTU level (Fig. 1). Nonetheless, no significant difference between NG and VG was seen.

**Figure 1.**
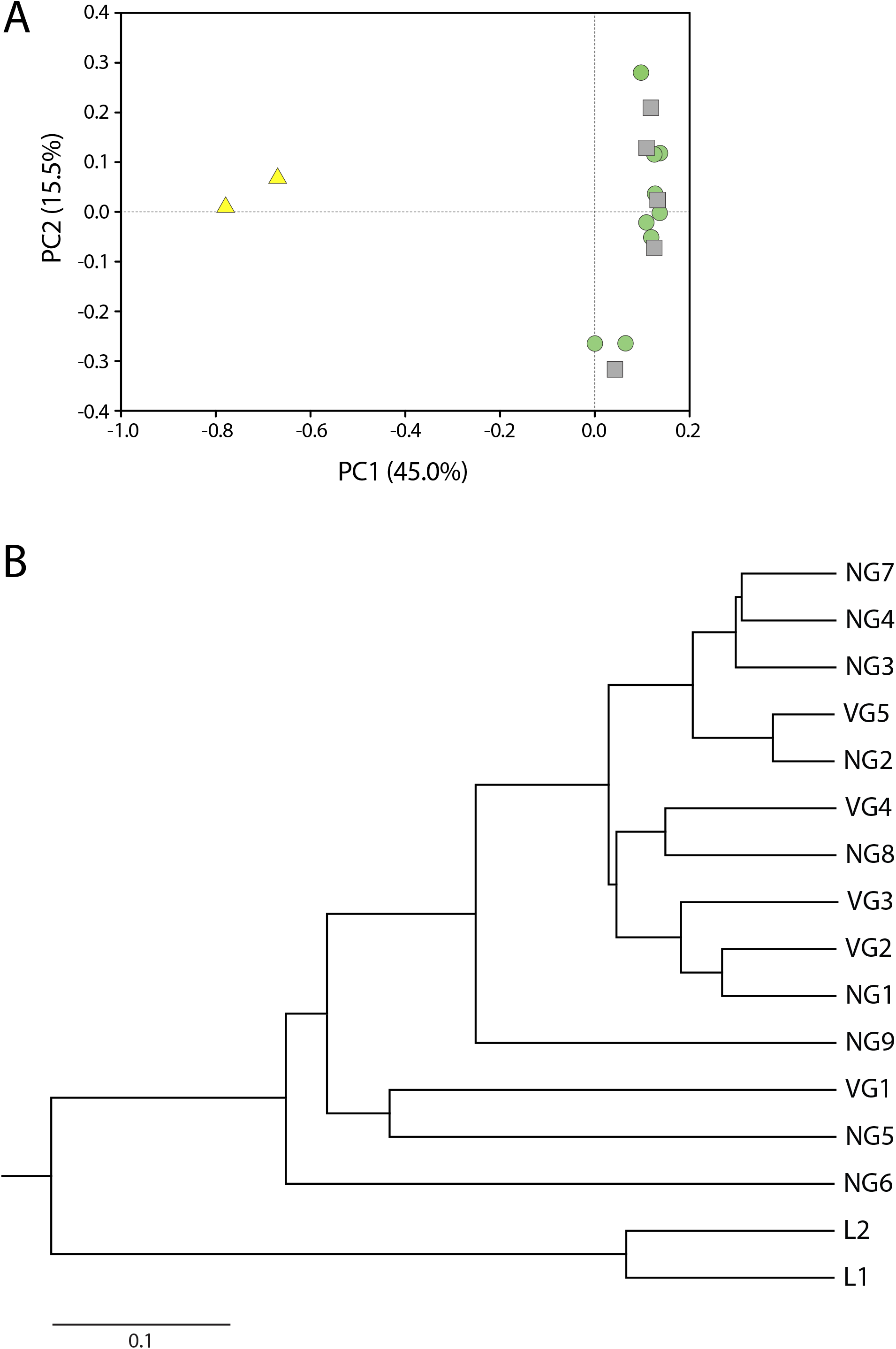
The relationships between the bacterial community profiles of the larva and adult bees, represented by a principal coordinates analysis (PCoA) plot (a) and an unweighted pair group method with arithmetic mean (UPGMA) clustering tree (b), based on Yue-Clayton dissimilarity metrics. The principal axes are shown with the percentage of variation explained between brackets. Each bee samples are denoted by larva (L, triangle, light yellow), non-varroa (NG, circle, light green), and varroa group (VG, square, light gray), respectively.

### The profiles of honeybee gut bacterial community

Although the alpha-diversity analysis indicated no discernible differences (i.e., diversity indices, Table 1), microbial communities were considerably different according to development stage (i.e., larva and adult) (Figs. 2-3).

**Figure 2.**
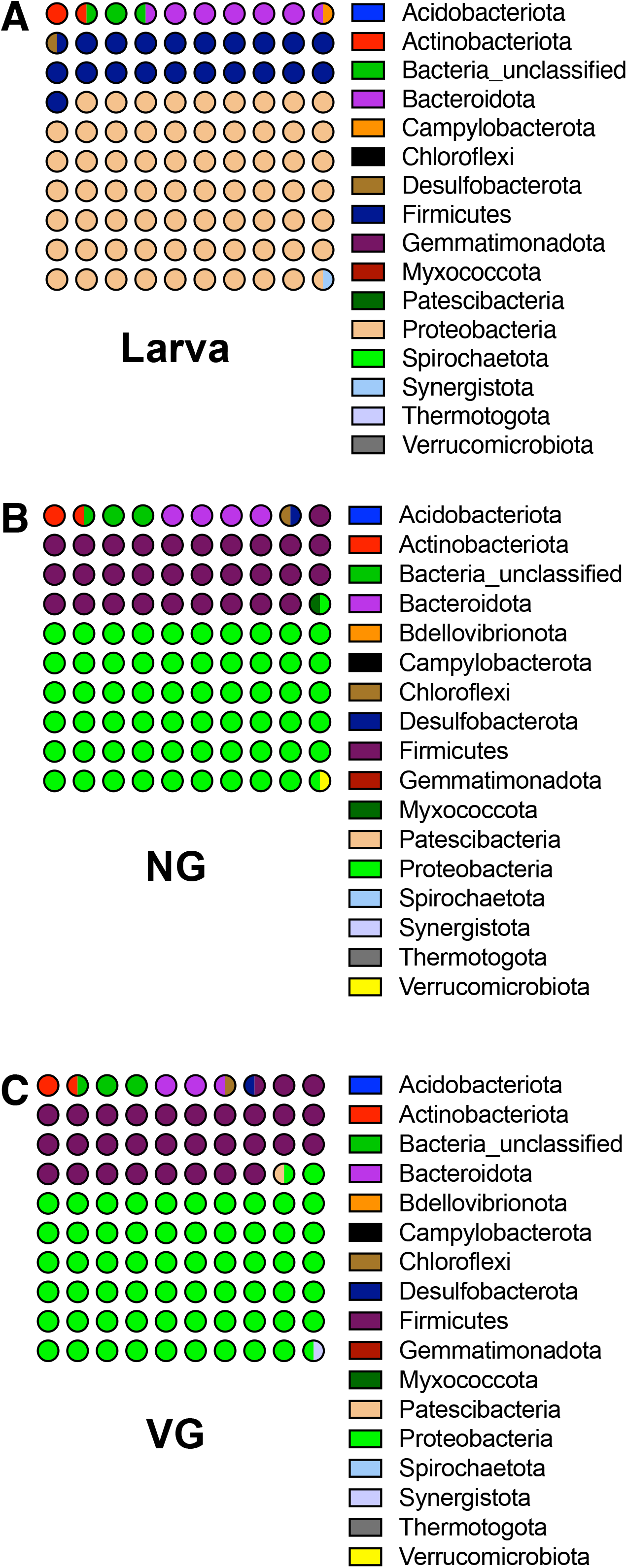
The relative abundances of the identified phyla in the L (a), NG (b), and VG (c) samples. Phyla abundances of taxa found in the difference groups represent by dot plot (10 × 10). Read sequences were assigned using mothur package and a reference database of recently updated 16S rRNA gene obtained from the Silva database (version silva.nr_v138.1).

**Figure 3.**
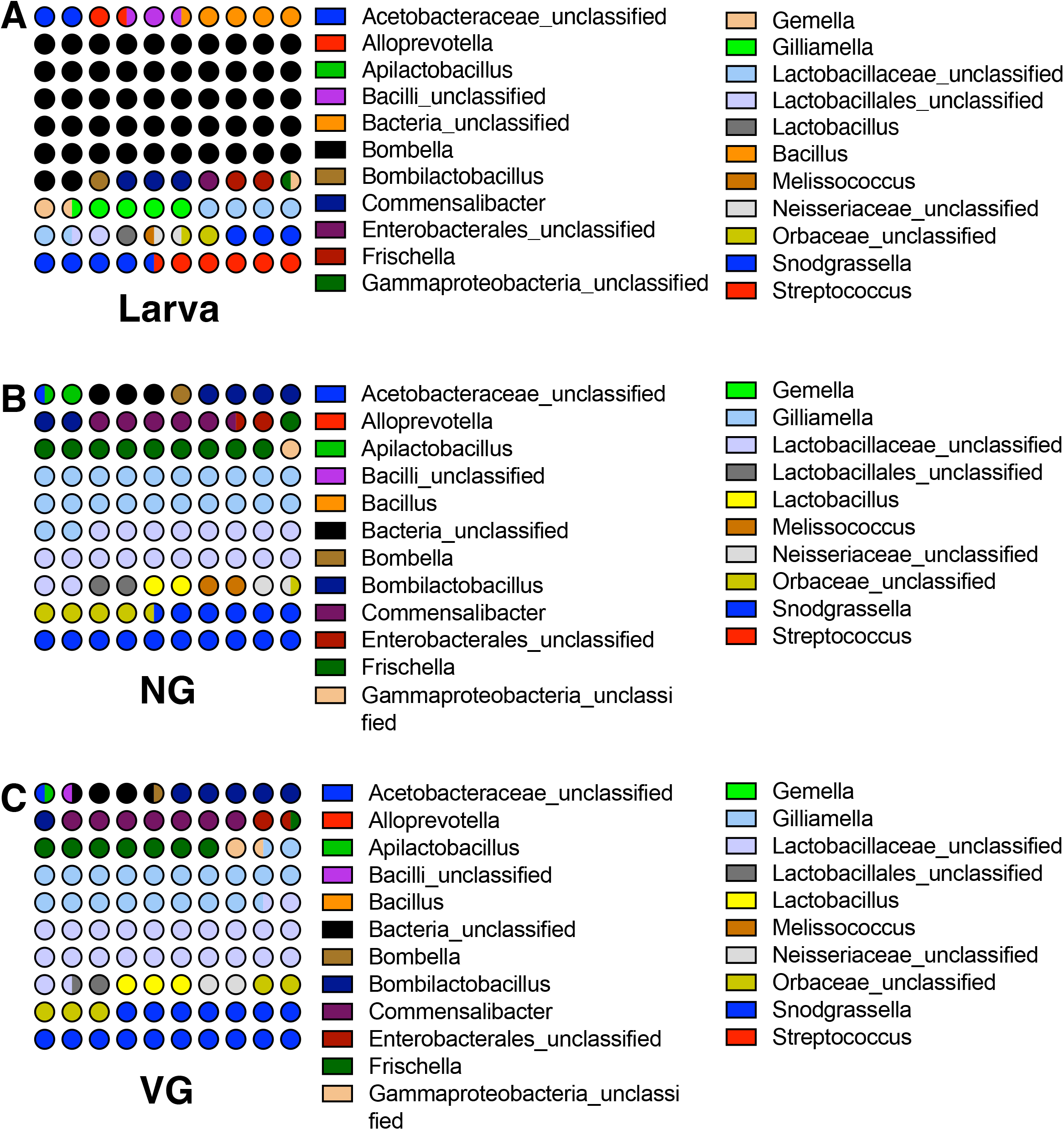
The abundances of the identified genera in the L (a), NG (b), and VG (c) samples. Genera abundances represent by dot plot (10 × 10). The selected most relatively dominated genera (more than 1% of total read sequences in each group) are shown in stacked. Read sequences were assigned using mothur package and a reference database of recently updated 16S rRNA gene obtained from the Silva database (version silva.nr_v138.1).

The analyzed sequence reads were classified into 40 phyla from all sample. We then estimated the relative abundances of a combined set of organisms collected by the same experimental sample (i.e., L group, NG, and VG). Finally, only 17 phyla were chosen as abundant phyla in this study (more than 0.1 percent of total reads in each sample) (Fig. 2). Proteobacteria (59.6-68.7 percent) and Firmicutes (20.4-30.7 percent) were identified as the most numerous phyla (>20 percent of total reads) in three groups (L group, NG and VG), followed by Bacteroidota, an unidentified group, and Actinobacteriota (more than 1% of total reads). In minor phyla (less than 1%), the phylum Campylobacrota was more abundant than both adult bee groups (NG and VG). On the other hand, Gemmatimonadota, Myxococcota, Synergistota, Verrucomicrobiota, and Bdellovibrionota were shown to be more abundant in NG than the L group and VG.

The analyzed sequence reads were classified into 727 genera at the genus level. Nonetheless, the majority of readings were classed as unclassified with a high taxonomic rank (i.e., family to class). We selected 22 genera from each group (threshold more than 1% of total reads) for sub-sequential analysis, including the unclassified group with a high taxonomic rank. Among the selected genera, we identified nine significant taxa (more than 4% of each group); *Bombella, Bombilactobacillus, Commensalibacter, Frischella, Gilliamella*, unclassified *Latobacillaceae*, unclassified *Orbaceae, Snodgrassella*, and *Streptococcus*. The genus *Bombella*, in particular, was found as a prominent species with the highest relative abundance in the L group (43.7 percent of total bacterial abundance). *Bombella* had a decreased the relative abundance in adult bees (less than 0.6 percent). Additionally, the L group had a higher the relative abundance of several taxa, including *Streptococcus, Alloprevotella, Bacillus, Gemella*, and other high-taxonomic groups (e.g., *Acetobacteraceae* and *Bacilli*), than the adult bees group (Fig. 3 and sFig. S2).

On the other hand, in NG and VG, a dominant microbe was identified as *Gilliamella*, unidentified *Lactobacillaceae*, and *Snodgrassella* (ranges 12.8 to 19.7 percent). Additionally, unclassified Enterobacterales, unclassified Gammaproteobacteria, *Lactobacillus*, and unclassified *Neisseriaceae* were identified as taxa with a higher abundance in NG and VG than the L group. Except for the genera *Apilacetobacillus* and *Melissococcus* or *Lactobacillus*, the relative abundances of the other genera were comparable between NG and VG (Fig. 3).

The LEfSe analysis was used to separate the distinctive taxa (at the genus level) from the inter-group (Fig. 4). We found no significant difference between NG and VG in the analysis. In light of that finding, we attempted to quantify the precise changes in microorganisms between the L group and adult bees from NG and VG. Seven genera were significantly enriched in the L group, led by *Bombella* (5.44 LDA score, p=0.0001), *Streptococcus* (4.47 and 0.0001, LDS score and p value, respectively), unclassified bacterial group (4.44 and 0.0001), unclassified *Acetobacteraceae* (4.35 and 0.0001), *Bacillus* (4.31 and 0.0001), *Gemella* (4.11 and 0.008), and *Alloprevotella* (4.11 (4.05 and 0.0001). While *Gilliamella*, unclassified *Lactobacillaceae*, unclassified *Orbaceae*, unclassified *Neisseriaceae, Frischella, Lactobacillus*, unclassified Gammaproteobacteria, *Bombilactobacillus*, and unclassified Enterobacterales were remarkably dominant genera in the adult bee group with a 4.43-4.93 LDA score and p<0.04. Interestingly, two genera, *Commensalibacter* and *Snodgrassella*, were identified as the major taxon in adult bees (NG and VG) (Fig. 3); nevertheless, the LEfSe analysis revealed no significant difference between the L group and adult bees (Fig. 4).

**Figure 4.**
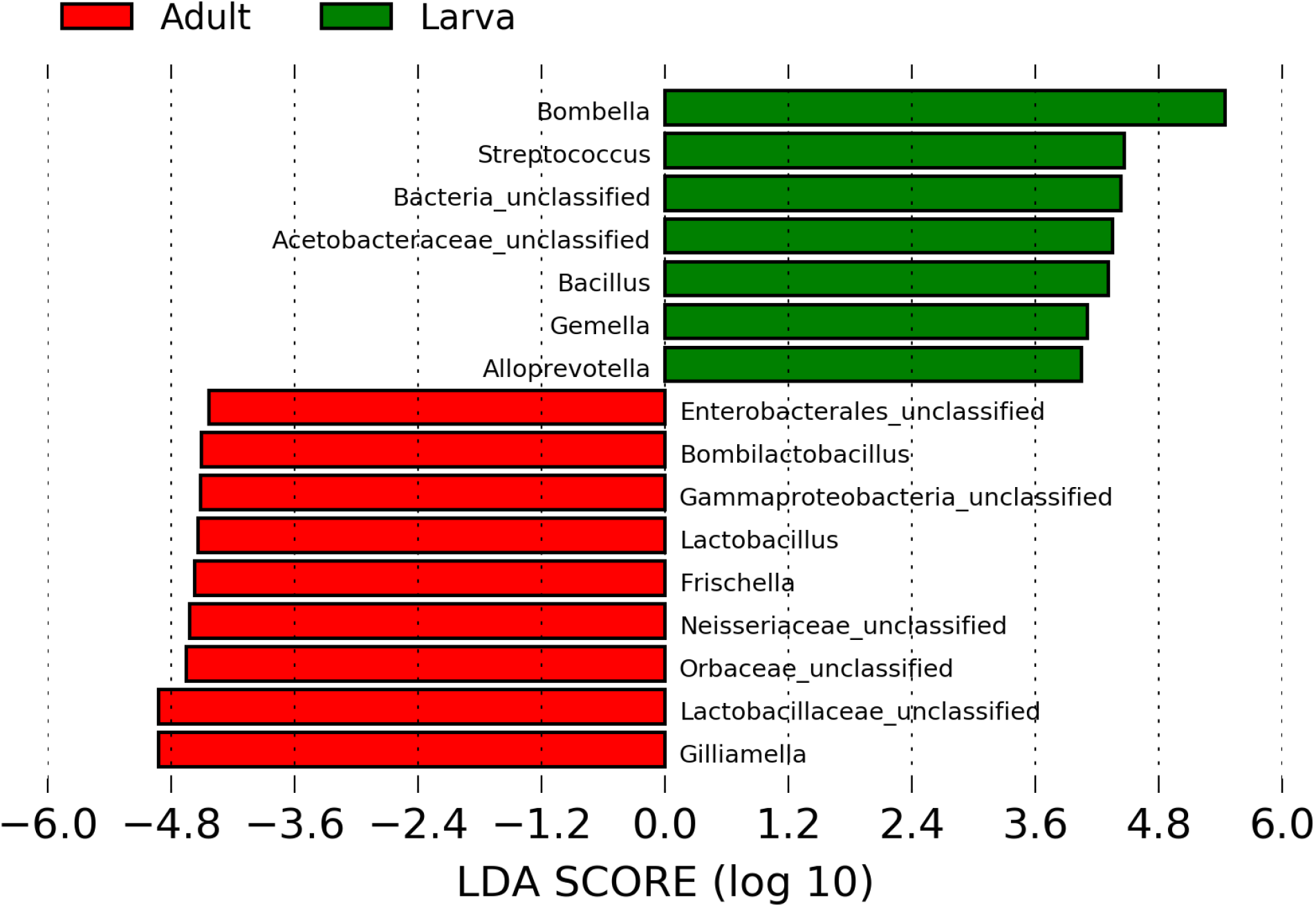
The LEfSe was carried out by Galaxy Project and the LDS score were presented by in the bar charts. LDA scores showed significant bacterial difference between Larva and adult bees (NG and VG) groups at the selected genera. The groups were statistically significant compared to each other (LDA > 2.0 and *p* < 0.05).

### Honeybee gut archaeal community profiles

In the current work, we attempted to investigate the archaeal community profiles of honeybees, including larvae. Surprisingly, we learned relatively little about the archaeal community from our ten samples, which included one larva and nine adult bees (NG, n=5; VG, n=4). Surprisingly, the genus *Methanomassiliicoccus* (phylum Thermoplasmatota) was found as the sole dominant bacterium in larvae. *Methanimicrococcus* (80.5 percent of total reads in NG) and *Candidatus* Methanoplasma (17.3 percent) were the most common microbes in NG, belonging to the Halobacteria and Thermoplasmatota, respectively. Additionally, an unidentified archaeal group was discovered in NG with a 2.0 percent abundance. When NG and VG were compared, *Methanomassiliicoccus* was a surprisingly dominant genus (98.7 percent of total reads in VG) (data not shown).

### Predicted functional profiles from bacterial communities

In general, it may be challenging to comprehend and extrapolate functional roles from the organization of microbial communities as determined by the 16S rRNA gene. As a result, putative functional profiles were inferred for inter-group comparisons using PICRUSt analysis and KEGG pathway information. The PICRUSt analysis revealed that the L group, NG, and VG all had enriched functional profiles in the bacterial population. The result of KEGG functional classes (levels 1 and 2) revealed substantial differences between the L and NG or VG groups in terms of functional categories (Fig. 5). Nonetheless, no significant variations in PICRUSt analysis were seen between NG and VG, similar to the results for alpha-diversity and LEfSe (Table 1 and Fig. 4).

**Figure 5.**
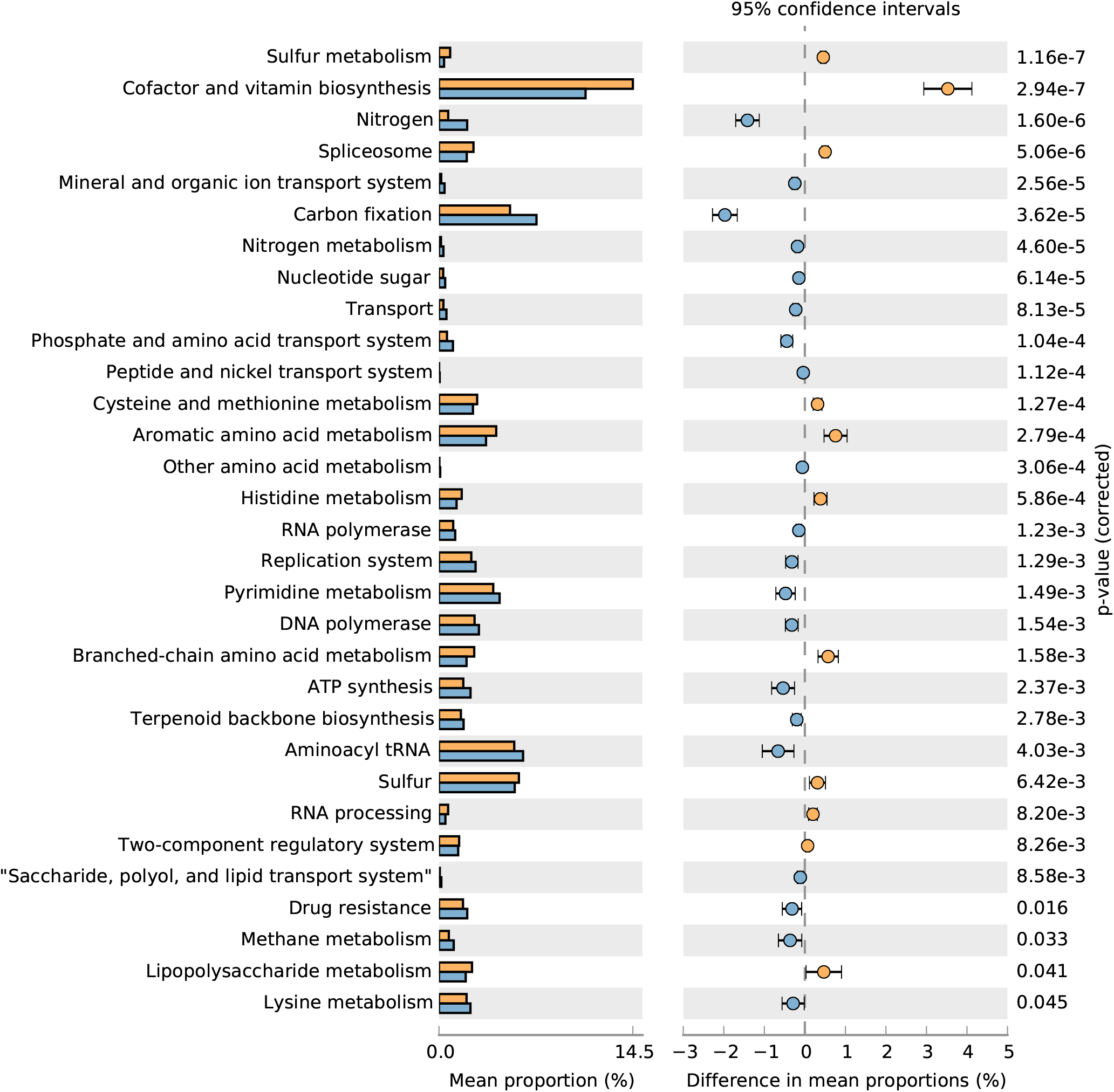
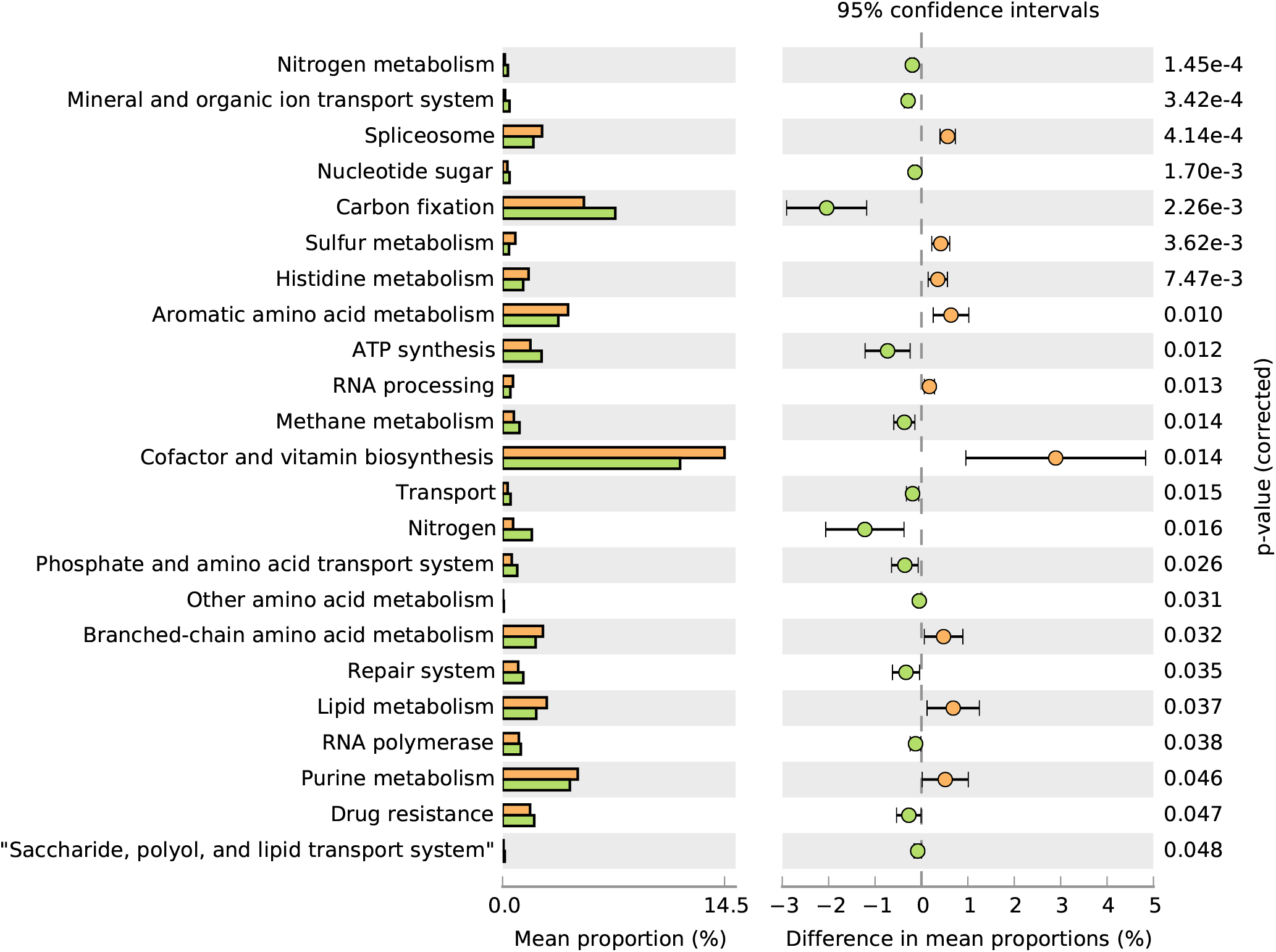
PICRUSt analysis. The chart for the predicted functional characterization at KEGG level 3 significant difference (*p* < 0.05) between larva and NG (a) or VG (b) groups was presented by STAMP software. Larva (orange), non-varroa group (NG, blue), varroa group (VG, green).

In comparison to NG or VG, the eight categories (lipid metabolism, splicesome, sulfur metabolism, cofactor and vitamin biosynthesis, RNA processing, histidine metabolism, aromatic amino acid metabolism, and branched-chain amino acid metabolism) had a significant effect size in the L group (effect size ranged from 0.37 to 0.82). In particular, as compared to adult bees, biotin synthesis gene clusters were identified in the L group (Fig. 5). Carbon fixation, methane metabolism, mineral and organic ion transport systems, nitrogen metabolism, glycosaminoglycan metabolism, nitrogen, nucleotide sugar, repair system, transport, peptide and nickel transport systems, and phosphate and amino acid transport systems all had a smaller effect size in the L group.

## Discussion

It has been demonstrated that the gut microbiota has a substantial functional role in organisms, including plants. Recently, the total number of microbial cells (i.e., bacteria) was recalculated to be comparable to the number of human cells, after previously being estimated to be at least tenfold (37). Regardless of the ratio, gut microbiota can have a substantial impact on host health and disease. Simultaneously, it was discovered lately that invertebrates, including bees, have a key link with a complex microbial community. Nonetheless, the critical physiological roles of the (bee) microbiota during the health or development stages are relatively unknown (24, 26). The use of NGS sequencing and culture-dependent techniques has considerably expanded our understanding of the link between bees and related bacteria, and the roles of gut microbiota have been recognized in the gut microbiota of the majority of healthy-adult worker honeybees.

The purpose of this study was to examine whether there were significant differences in the gut microbiota of *Varroa*-infected (attached) bees and those were not infected. Additionally, the complete larval body’s microbiota was employed to create a separate group. To address these questions, we sequenced the 16S rRNA gene and analyzed archaeal and bacterial populations from the samples using the Illumina NovaSeq technology.

As previously reported, Bacteroidetes and Firmicutes are the predominant taxa for the vertebrate gut microbiota (references in 36). However, as previously documented in several other studies on honeybees, we discovered that larva and adult groups were dominated by the Proteobacteria and Firmicutes phyla (38-42). We identified the main phyla and genera in each experimental group and discovered substantial differences in the bacterial community between the L and adult bee groups (combined with NG and VG). However, as previously stated, biostatistical analysis indicates no difference in the resultant alpha diversity and bacterial community structure between NG and VG. Hubert et al. reported that various microorganisms (e.g., *Morgnaella, Sprioplasma*, and *Arsenophonus*) were transmitted between honeybees and *Varroa* (34). Additionally, Gammaproteobacteria, Betaproteobacteria, *Lactobacillus* spp., *Bifidobacterium* spp., and *Brevibasillus laterosporus* were detected in adult bees from *Varroa*-infested colonies using the qPCR technique (35). Thus, our hypothesis regarding sampling time is explicable; in this study, we collected samples from hives treated with anti-*Varroa destructor* medicine. However, the two studies cited previously (34, 35) evaluated the total body, not only the gut flora.

Interestingly, the phylum Bacteroidota (formerly termed Bacteroidetes) had a larger relative abundance (2.16-5.75 percent) than Actiobacteriota (1.28-1.62 percent) in all samples (Fig. 2), as previously described (40, 43). Particularly, Bacteroidota in L group was more abundant than adult bees. However, the Bacteroidota abundance observed in adult bees was higher than that of the other studies (39, 41, 42). The variables underlying these discrepancies are likely to be experimental differences, such as the target region for the 16S rRNA gene (e.g., V1-V3 or V3-V4) (44).

Bacteroidota is a prominent taxon in both mammalian and insect gut microbiota (45-47). The phylum Bacteroidota can degrade and utilize soluble polysaccharides via polysaccharide utilization loci-like systems (46). It is obvious that extracellular enzymes from bacterial cells of the phylum Bacteroidota can contribute to supply vitamins to host through intra-or intercellular reaction chains (48). However, due to the low abundance of the phylum Bacteroidota, its functional roles in honeybees are less understood than those of other taxa such as Firmicutes (26).

Similarly to other previous studies, we discovered distinct genera in both groups at the genus taxon level (larvae and adult bees). Particularly, in L group, the genus *Bombella*-related reads were dominant. Li et al. first proposed the genus *Bombella* as a member of the Alphaproteobacteria (49), with the description of *Bombella intestine* as the type species from bumble bee crop. At the time of this writing, the genus *Bombella* has just four validly named species (49-51). Interestingly, these *Bombella* spp. have been only isolated from honeybee-associated environments such as honeycombs and gut. Also, all members of the genus *Bombella* share unique characteristic for acetic acid producing. Indeed, It has been already predicted that *Acetobacteraceae* Alpha 2.2 bacteria of the genus *Bombella* plays an functional roles on the young larval fitness (52). Additionally, recent genomic studies clearly show that they play a critical role in the interaction with their host (53, 54). The genus *Allopreovotella* was only identified as a minor group inside the L group, among several other genera and high-taxonomic groups (e.g., *Acetobacteraceae* and Bacilli) (Fig. 3). Indeed, only one study has identified the genus *Alloprevotella* in adult bees to our knowledge (39). The genus *Alloprevotella* reclassified from *Prevotella* (55) can produce acetate and succinate as end products (i.e., short-chain fatty acid, SCFA) from glucose via fermentation. The SCFAs are the main metabolites produced by gut bacterial fermentation using saccharides (e.g., starch or fiber) and contribute to crucial physiological effects on host health including immunity, behavior, or neurological disorders (56, 57). In fact, the roles of the *Prevotella* spp. in the host are uncertain and various investigations have reported contradictory interpretations (58). Nonetheless, it may be difficult to dismiss *Alloprevotella* as a beneficial commensal because it plays a critical role in larval health via polysaccharide breakdown and SCFA production.

In comparison to the L group’s gut microbiota, adult bee groups contain a distinct microbial community constituted of four distinct classes (e.g., Bacilli, Alpha-, Beta- and Gamma-proteobacteria) (Fig. 2). This could be due to the four distinct developmental stages (egg, larva, pupa, and adult) and the nursing bees’ distinct-yet-simple feeding systems for larvae (pollen and honey) (26, 59). These four classes have been given the names *Bombilactobacillus, Commensalibacter, Frischella, Gilliamella, Lactobacillus*, and *Snodgrassella*, and their roles in honeybees have been thoroughly characterized (25, 27, 60, 61). Additionally, the genus *Bifidobacterium* is known as a core bacterial clade and may provide organic or aromatic compounds degraded from pollen (62). These pollen-derived compounds might be cross-fed to other gut bacterial members and finally contribute to bee development (63). Despite the fact that there are only two studies linking microbiota and *Varroa* infection, the results indicate that *Bifidobacterium* was detected in gut microbiota and its abundance was positively correlated with *Varroa* infection (33, 35). Unexpectedly, in the present study, the relative abundance for *Bifidobacterium* (Actinobacter phylum) were less than 0.5% of total microbiota in both L and adult bee groups. However, the relative abundance of unclassified *Bifidobacteriaceae* as a high taxon level was similar to the genus *Bifidobacterium*. This indicates that the honeybee gut has a greater number of unclassified species belonging to the family *Bifidobacteriaceae*. Lactic acid bacteria such as *Bifidobacterium* protect hosts from (opportunity) pathogen infection by lowering the pH of the gut environment through the production of organic acids and antimicrobial substances such as antimicrobial peptides (AMPs) (64). As a result, it is worth mentioning that clarifying their potential activities may aid in our knowledge of the honeybee gut microbiota’s more functional roles in pathogen defense.

In the present study, we firstly attempted to analyze and identify archaeal community in honeybee gut. However, unexpectedly, the archaeal diversity and community structure was extremely limited. Only a few bee samples harbored methanogen, despite the honeybee gut was anoxic condition with partial oxygen pressure closed to zero (65). This could be because of the positive redox potential (215-370mV) (65). Under anaerobic conditions with a negative redox potential, methanogenesis is conceivable (−200mV) (66, 67). Indeed, only a few insects, including beetles, cockroaches, termites, and millipedes, possess methanogen or other archaeal groups in their hindguts (45, 68).

This study has the following limitations: i) gut microbiota analysis was performed a few days following therapy for *Varroa* infection. Examining the time course of *Varroa* infection on the honeybee gut may provide more detailed information about microbiota changes (i.e., dysbiosis); ii) the sample size for each group was relatively small (see materials and methods), despite the fact that our results were consistent with previous studies. Nonetheless, this study shows that the organization of the honeybees’ bacterial community reflects their developmental stage. The L or adult bees group has a straightforward and distinctive bacterial composition and distribution of the several bacterial groups. The expected functional profiles of L and adult bee groups are distinct depending on their bacterial communities. These functional characteristics, however, were comparable across NG and VG. Finally, the bacterial and archaeal community structure seen in honeybees may be an essential factor in the health of the bees.

### Outlook

To date, most investigations have focused on pathogenic or beneficial microorganisms in human gut, which have an influence gastric diseases, metabolic syndrome, brain, or behavior (69-71). However, it makes difficult to fully characterize the microbial composition and to isolate the key microorganisms from the microbiota. Numerous microorganisms remain uncultured, and the impacts of specific microorganisms cannot be examined using molecular techniques (i.e., NGS). Additionally, direct study on the relationship between humans and microbiota or its physiological involvement in the host gut is restricted. Invertebrate organisms (i.e., insect) harbor relatively simple gut microbial communities (45). Beyond the fruit fly *Drosophila* (72), honeybees, in particular, have been used as a model organism to study social behavior, brain disorders, aging, and development (73-76). The honeybee’s microbial (i.e., bacterial) community structure is straightforward, and only a few bacterial species have been identified as being prevalent in the gut system, which is another advantage of using honeybees as a model for gut microbiota (25, 26).

## Acknowledgements

We are grateful to Greenbees Co., (http://greenbees.kr/) located on Jeju Island, for providing us with honeybee samples and for engaging us in a fruitful conversation about the effects of varroa infection.

## Funding

This work was supported by grants from the National Research Foundation of Korea (No. 2020R1I1A3062110) and Startup funds of HIT Center for Life Sciences.

## Author contributions

WJK and SJP designed the experiments. MK and SJP performed the experiments. WJK provided support for the experiments. MK, WJK, and SJP analyzed the data. WJK and SJP wrote the manuscript. All authors read and approved the final manuscript.

## Availability of data and materials

The raw reads recovered in this study were deposited in the DDBJ/ENA/GenBank Sequence Read Archive (SRA) under the study accession number PRJNA823814.

## Code availability

Not applicable

## Declarations

### Conflicts of interest

The authors declare that there are no conflicts of interest.

### Ethics approval

Not applicable

## Supplementary figure legends

**Figure S1**. Alpha-diversity, estimated by Chao1 (a), Shannon richness (b), and inverse Simpson diversity (c) indices is plotted for larva (L, light yellow), non-varroa (NG, light green), and varroa group (VG, light gray). These values were calculated with Yue and Clayton dissimilarity metric based on the proportions of OTUs in different samples. The plots are based on the data shown in Table 1. The horizontal line inside the box indicates the median value. The whiskers represent the lowest and highest values within the 1.5 times interquartile range (IQR) from 25^th^ and 75^th^ percentile. Outliers as well as individual sample values are shown as dots.

**Figure S2**. Heatmap showing the microbial taxa selected most relatively dominated genera (more than 1% of total read sequences) in each group. Heatmap was generated using gplot package.

## Notes

### Competing Interest Statement

The authors have declared no competing interest.

