## Supplementary figures 1-2 for "Gut microbiota analysis of the western honeybee (*Apis mellifera* L.) infested with the mite *Varroa destructor* reveals altered bacterial and archaeal community"

Supplementary Figure S1a.

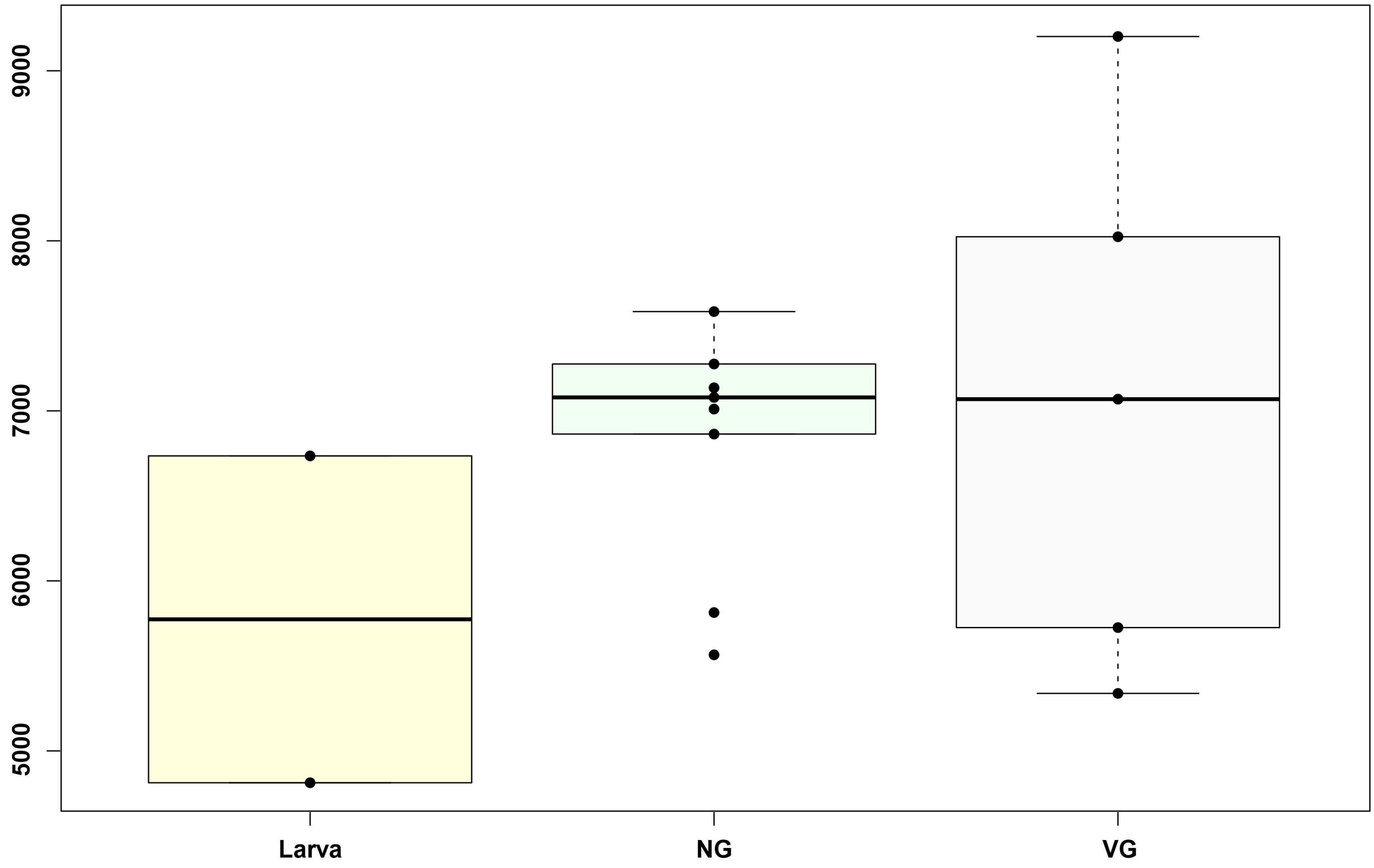

Supplementary Figure S1b.

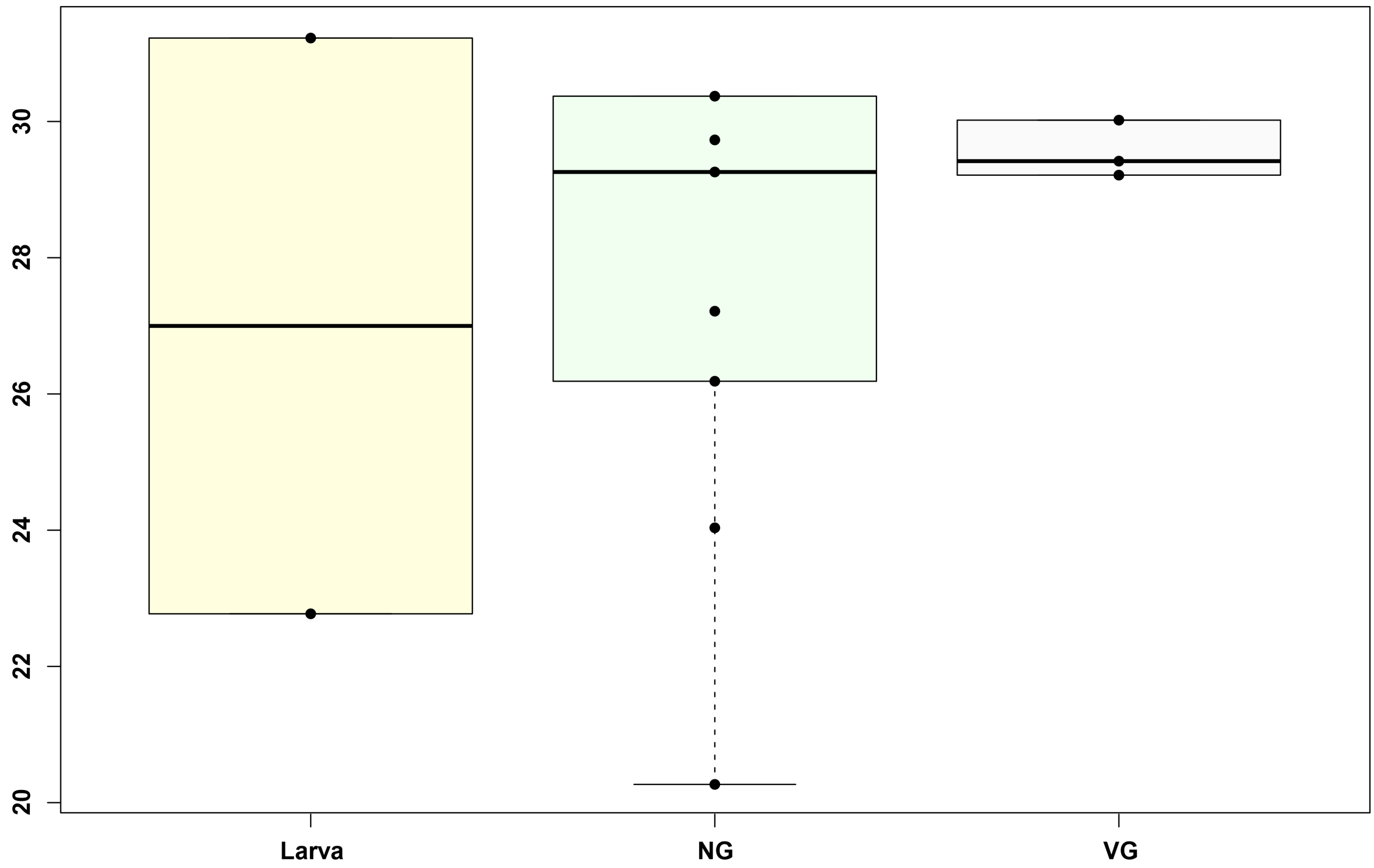

Supplementary Figure S1c.

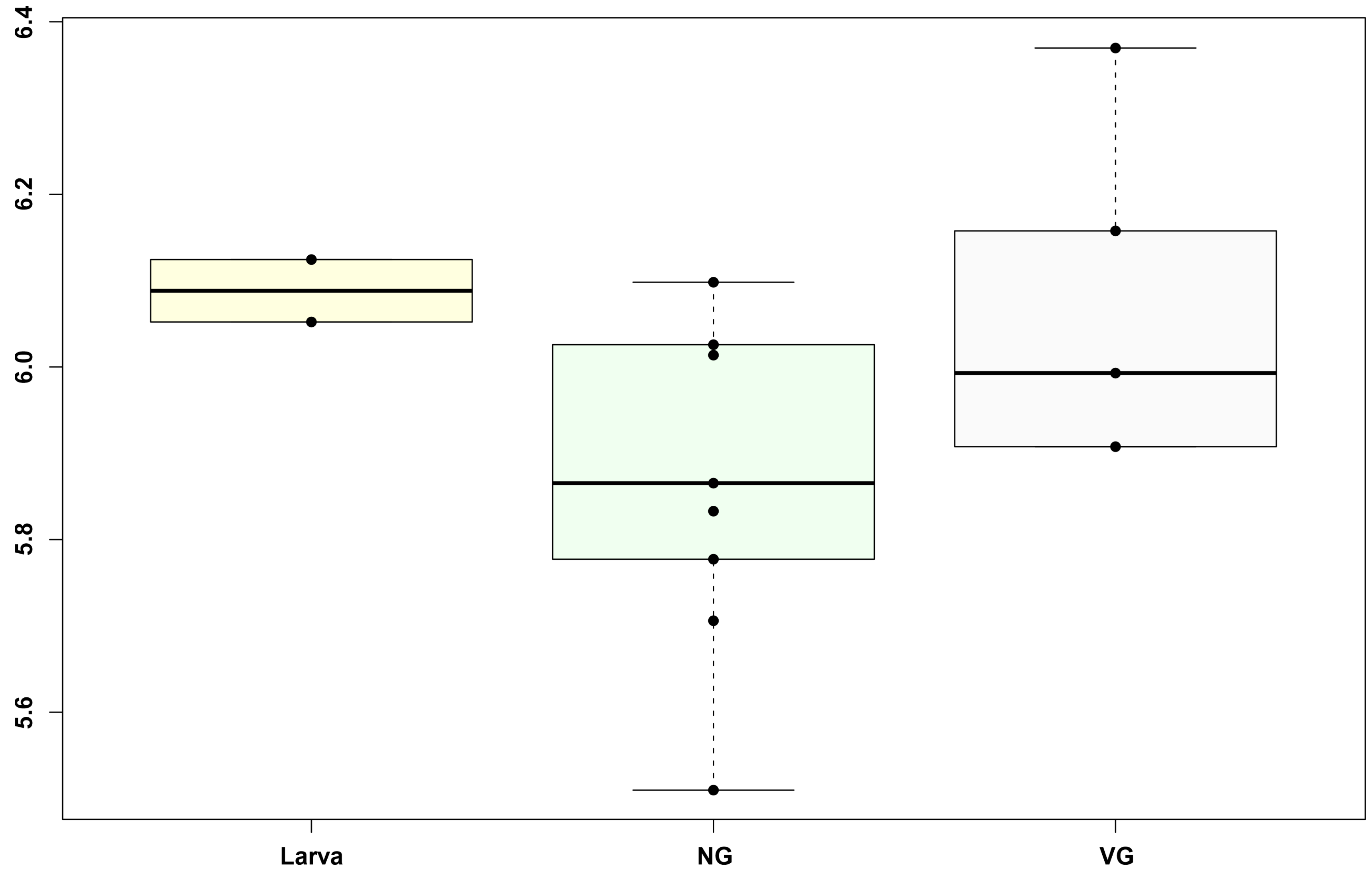

### Supplementary Figure S2.

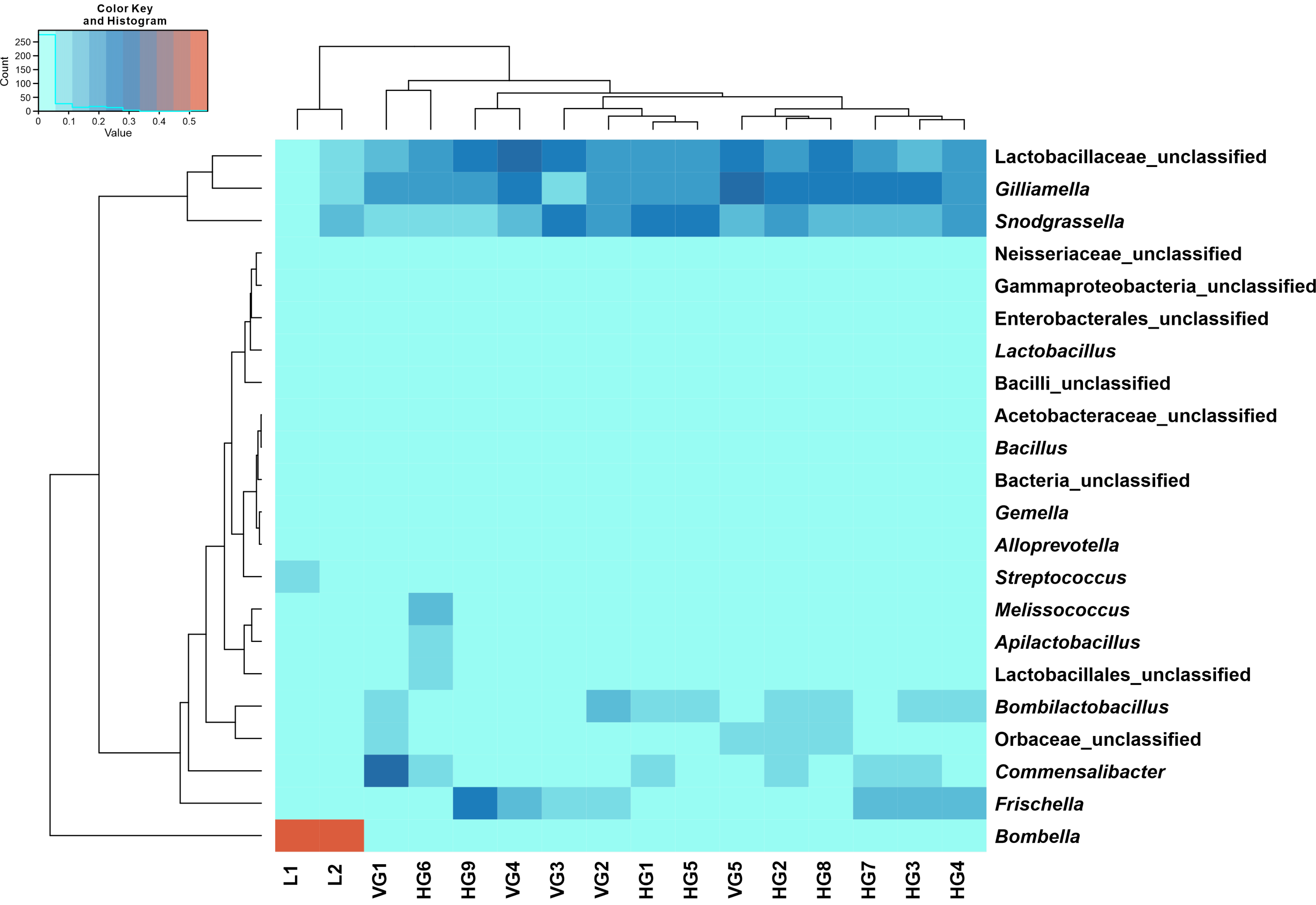
